# Rates of evolution differ between cell types identified by single-cell RNAseq in threespine stickleback

**DOI:** 10.1101/2024.12.06.627160

**Authors:** Maria L. Rodgers, Swapna Subramanian, Lauren E. Fuess, Wan He, Samuel V. Scarpino, Andrea J. Roth-Monzón, Daniel Jeffries, Martine Seignon, Kathryn Milligan-McClellan, Rebecca Carrier, Natalie C. Steinel, Daniel I. Bolnick

**Affiliations:** University of Connecticut, Department of Ecology and Evolutionary Biology, Storrs, CT, 06269; North Carolina State University, Center for Marine Sciences and Technology, Department of Biological Sciences, Morehead City, NC 28557; Texas State University, Department of Biology, San Marcos, TX 78666; Institute of Experiential Artificial Intelligence, Northeastern University, Boston MA 02115; Division of Evolutionary Ecology, Institute of Ecology and Evolution, University of Bern, Bern, Switzerland; The Jackson Laboratory for Genomic Medicine, Farmington, CT, USA, 06032; University of Connecticut, Department of Molecular and Cell Biology, Storrs, CT, 06269; Department of Chemical Engineering, Northeastern University, Boston MA 02115; University of Massachusetts Lowell, Department of Biological Sciences, Lowell, MA, 01854

**Keywords:** evolutionary rates, *Gasterosteus aculeatus*, single cell sequencing, immunity

## Abstract

Rates of evolutionary change vary by gene. While some broad gene categories are highly conserved with little divergence over time, others undergo continuous selection pressure and are highly divergent. Here, we combine single-cell RNA sequencing (scRNAseq) with evolutionary genomics to understand whether certain cell types exhibit faster evolutionary divergence (using their characteristic genes), than other types of cells. Merging scRNAseq with population genomic data, we show that cell types differ in the rate at which their characteristic genes evolve, as measured by allele frequency divergence among many populations (F_ST_) and between species (dN/dS ratios). Neutrophils, B cells, and fibroblasts exhibit elevated F_ST_ at characteristic genes, while eosinophils in the intestine and thrombocytes in the head kidney exhibit lower F_ST_ than the average for 1000 random genes. Gene network centrality also differed between immune- and non-immune-associated genes, and closeness centrality was positively related to gene F_ST_. These results highlight the value of merging single cell RNA sequencing technology with evolutionary population genomic data, and reveal that genes which define immune cell types exhibit especially rapid evolution.

## Main text

Comparative evolutionary genetics studies have clearly established that the rate of evolutionary change varies between genes^1–3^. Some categories of genes are highly conserved and exhibit very slow evolutionary divergence between populations or species indicating negative selection purging new mutations; other genes exhibit a signature of persistent positive selection favoring new mutations. For example, genomic divergence among human populations is, on average, especially rapid for immune genes, particularly those involved in anti-helminth immunity^4^.

However, such studies tend to use coarse gene ontology categorizations to classify genes, which may not effectively reflect the function of, or relations between, genes. This limitation is especially acute for non-model organisms whose gene ontologies are inferred from very distantly-related models. An alternative approach would be to fuse the relatively new technology of single-cell RNA sequencing (scRNAseq) with evolutionary genomics. In non-model organisms, scRNAseq can provide a data-driven tool for assigning sets of genes to the cells in which they function. This allows us to ask, for the first time, whether certain types of cells exhibit consistently faster evolutionary divergence (at their characteristic genes), than other types of cells.

Here, we apply this approach to ask whether different cell types exhibit consistently different rates of molecular evolution. To define cell types we use a multi-tissue scRNAseq dataset from threespine stickleback (*Gasterosteus aculeatus*), an emerging model organism in evolutionary genetics. We identified clusters of cells with similar transcriptome profiles, and a set of genes whose expression is diagnostic for each cluster (cell-type specific genes, CTSGs). We then used a large population genomic dataset to identify the extent of among-population allele frequency divergence at each of the CTSG loci. We also measure signatures of selection in between-species divergence. For comparison to these CTSGs, we also evaluate evolutionary rates for a set of randomly chosen loci that are not cell-type-specific. We show that the rate of CTSG evolution varies among cell types, and is faster for some cell types involved in immunological processes. This variation in evolutionary rates is in part associated with cell differences in gene network architecture.

We obtained scRNAseq data from upper intestines, lower intestines, liver, and gills (N = 3 fish each (note: the same 3 fish were used for all organs)) and head kidneys (a.k.a. pronephros, 9 fish total) (see Table S1 for details). Libraries for intestinal tissues ranged from 12,278-30,789 estimated cells (mean reads per cell ranged from 13,173-35,513), head kidneys ranged from 8,161-19,382 estimated cells (mean reads per cell ranged from 15,580-55,204), livers ranged from 8,436-9,662 estimated cells (mean reads per cell ranged from 32,088-52,690), and for gills estimated cells ranged from 7,327-9,089 (mean reads per cell ranged from 41,759-65,325). For intestines, median genes per cell ranged from 257-779, in the head kidneys, median genes per cell ranged from 307-707, in livers median genes per cell ranged from 309-372, and in gills median genes per cell ranged from 327-380.

t-SNE clustering analysis identified 23 clusters of cells for the intestine (24 in the head kidney, 4 in the liver, 5 in the gills). Fuess et al previously characterized 8 distinct cell clusters in the stickleback head kidney^5^. We were able to clearly assign a known cell type to 11 of the intestinal cell clusters, based on the identity of genes with elevated expression within a given cluster (Fig. 1a). All clusters in the liver and gills were able to be identified. Each cell cluster was diagnosed by high expression of at least one gene (typically multiple genes), which we refer to as cell-type specific genes (CTSGs). Some examples of CTSGs from the intestine are plotted in Figure 1.

**Figure 1.**
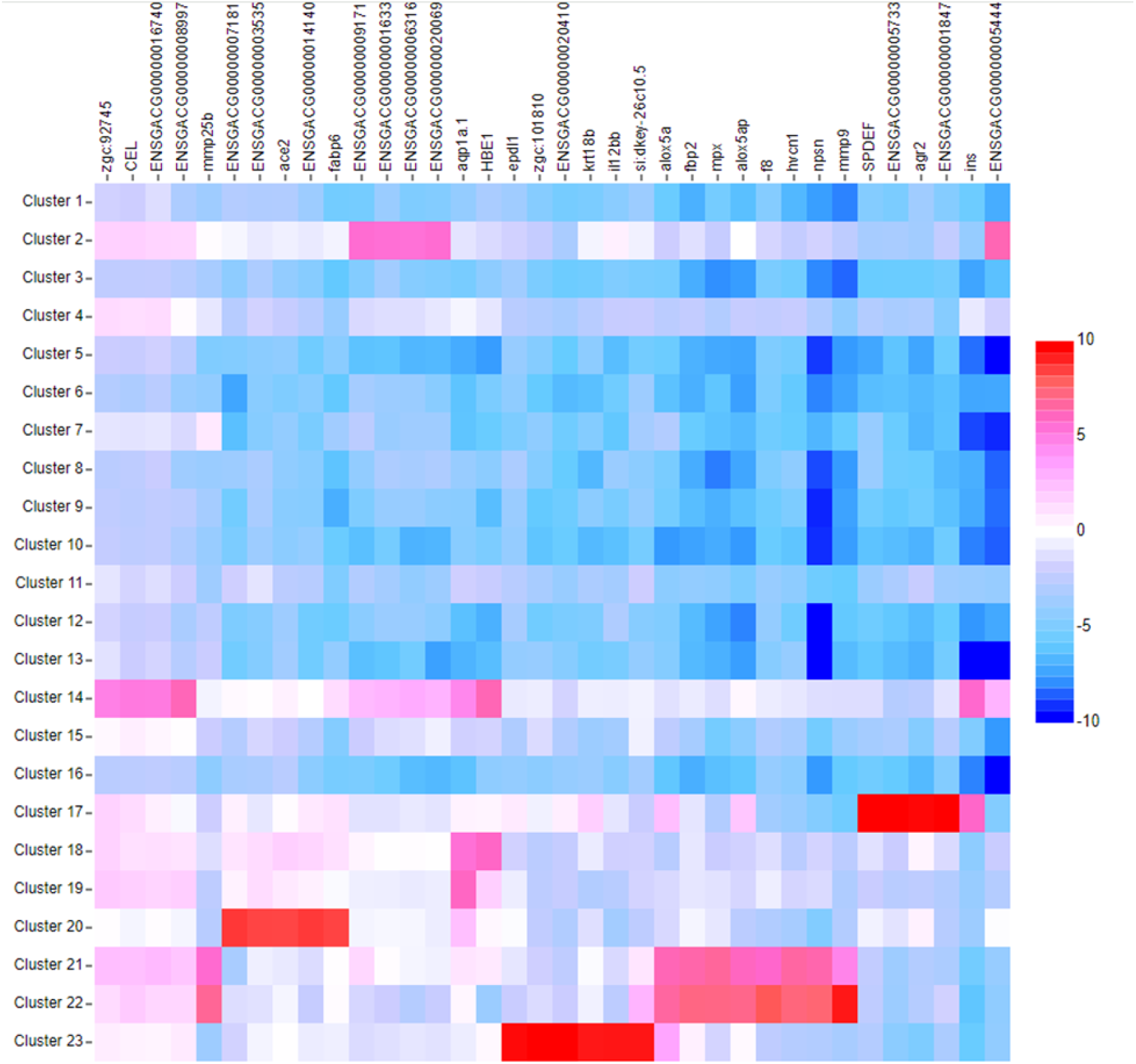
Heatmap of the intestinal cell clusters. Rows are cell clusters and gene transcripts are columns (with either gene name if annotated in the reference genome, or transcript ID). Red cells represent genes whose expression is especially high in a particular cell cluster, thus identifying cell type specific genes (CTSGs) whose evolutionary rates we query. This figure was made using Loupe Browser 7.

For instance, *cel* is a gene expressed at high frequencies only in cell cluster 14, and in humans is transcribed in the pancreas and in the mammary gland during lactation, allowing us to define this cell cluster as acinar cells^6^. Cluster 17 is identified as intestinal epithelial cells based on strong expression of several genes, including *spdef* and *agr2*, two genes strongly associated with goblet cells (a specialized type of epithelial cell)^7^. The 11 identified clusters in the intestine are: hematopoietic cells, stem cells, eosinophils, acinar cells, intestinal epithelial cells, erythrocytes, proerythroblasts, enterocytes, neutrophils, myeloblasts, and monocytes (Supplemetary Table 1). The four identified clusters in the liver are: neutrophils, erythrocytes, hepatocytes, and Kupffer cells (Supplementary Table 2). The five identified clusters in the gill are: erythrocytes, neutrophils, natural killer cells, B cells, and epithelial cells (Supplementary Table 3).

Next, we estimated rates of microevolutionary divergence (F_ST_) in stickleback using whole genome sequences from 28 stickleback populations (2 marine populations as an outgroup and from various-sized lakes in Alaska (N = 11) and on Vancouver Island (N = 15 lakes). From each population we pooled ~100 individuals’ DNA for PoolSeq to obtain genome-wide allele frequency estimates from each population (see Weber et al 2022 for details and population information)^8^. We then calculated pairwise F_ST_ to measure between-population allele frequency differences at each SNP in the genomic dataset, for all pairwise comparisons between populations. A phylogenetic tree confirms substantial population divergence among all lakes and between marine and freshwater populations (Fig. 2a), with Alaskan and Vancouver Island lakes forming distinct clades, and within-region genetic structure corresponding closely to different watersheds and sub-regions. For present purposes we focus on selection associated with the dramatic ecological shift when marine stickleback colonized freshwater habitats, so we focus on pairwise F_ST_ between an Alaskan anadromous marine population (Rabbit Slough) and 8 native Alaskan lake populations (excluding artificially founded lake populations), and F_ST_ between a Vancouver Island anadromous marine population (Sayward Estuary) and 13 Vancouver Island lake populations (excluding two suspected of being recent marine invasions). For each SNP we calculated the mean F_ST_ across all marine-freshwater contrasts. Next, we took the average of these mean F_ST_ values, across all SNPs within a given gene (including introns and a 5 kb window on either side) to obtain a gene-level measure of among-population divergence. Some loci exhibit fixed or nearly fixed differences in allele frequency indicative of strong divergent selection (Fig. 2b). A genome-wide scan of these averaged F_ST_ values (Fig. 2c) confirms that there is among-gene variation in the rate of evolutionary divergence.

**Figure 2:**
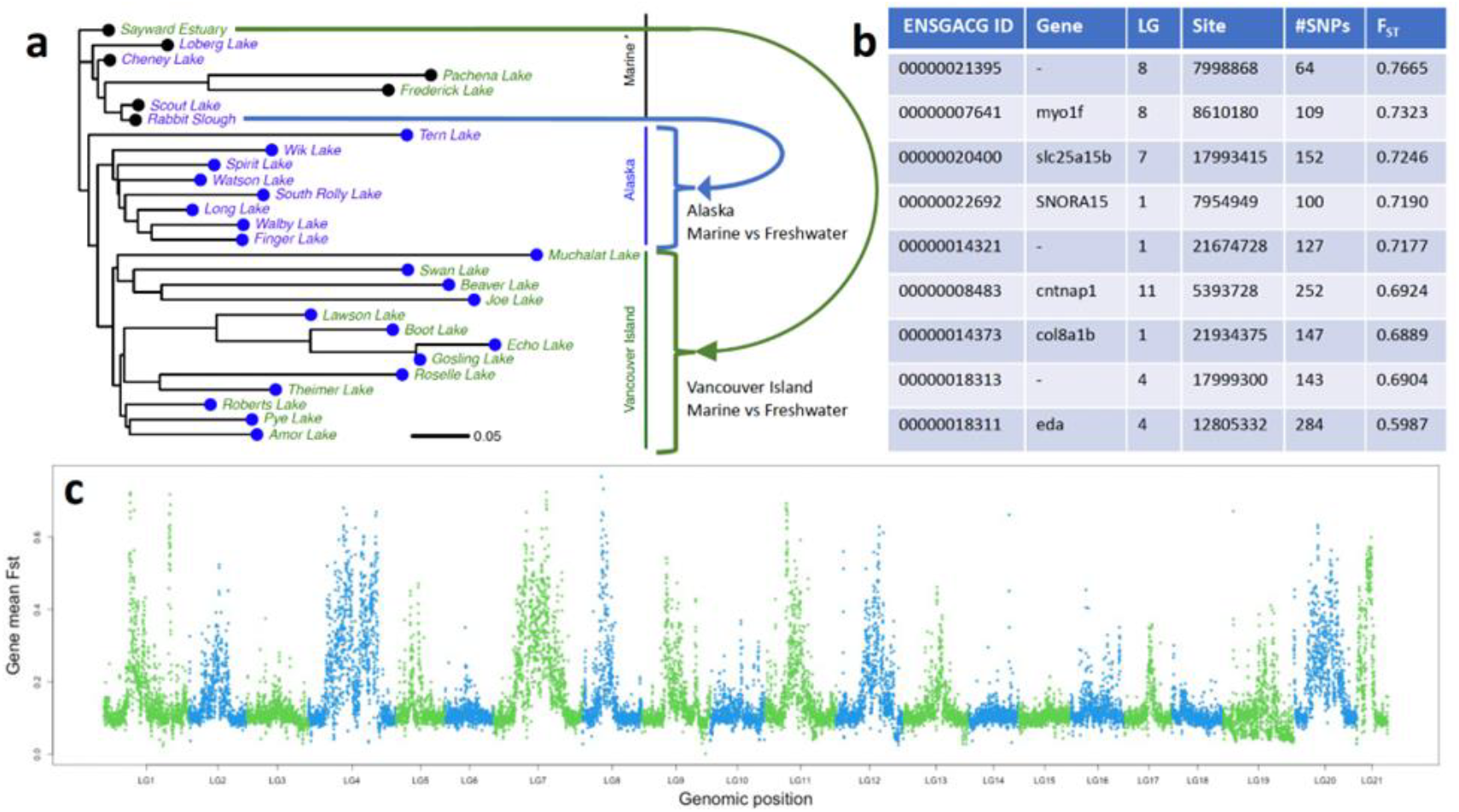
**a.** A phylogenetic tree of 28 stickleback populations based on pairwise Fst values for all Chromosome 15 SNPs identified with Poolseq. Freshwater populations are colored blue, marine populations are colored in black including three Alaskan lakes in which marine fish were introduced to lakes in the past few decades by humans (Loberg, Cheney, and Scout) and two lakes on Vancouver Island where a historical tsunami is believed to have caused a replacement with marine fish (Frederick and Pachena). Alaskan lakes form a clade, and Vancouver Island lakes form a clade (setting aside known or presumed marine introductions). We calculated marine to freshwater genetic divergence within each region, then averaged these values for each SNP, then averaged across SNPs within each gene (with flanking sequences. **b.** A table of 9 genes with the highest mean Fst, showing the ENSGACG gene ID, gene name (if known), linkage group, base position for the center of the gene, the number of SNPs within the gene and 5kb flanking region, and mean Fst across the SNPs. The 10th row is *eda*, a well-documented target of selection during marine to freshwater evolution. **c.** A Manhattan plot showing mean marine-freshwater Fst for each gene (points), coloring shows alternating linkage groups.

This among-gene variation in the strength of divergent selection is, in part, attributable to the cell type within which the genes function. Merging the scRNAseq results with our population genomic data, we find that selection (e.g., exceptionally high F_ST_) is, on average, stronger for CTSGs associated with certain cell types, than for other cell types for both intestinal cell clusters (Fig. 3a) and head kidney cell clusters (Fig. 3b). Using nested linear mixed models, we tested whether SNP F_ST_ values depended on the CTSG gene they are close to, with genes nested within cell types. Likelihood ratio comparisons of models with fixed effects of cell type versus just an intercept confirmed significant among-cell-type variation (log likelihood ratio LLR = 45.5, df = 17,4, P < 0.0001). Erythrocytes for example averaged F_ST_ of 0.106, compared to 0.270 for B cells. The cell types with elevated Fst at CTSGs include neutrophils, B cells, and fibroblasts (nested mixed model ANOVA, LLR = 37.16, df = 11,4, P < 0.0001). We also contrasted CTSG F_ST_ values against the gene-level mean F_ST_ for 1000 randomly selected genes from the genome. For instance, fibroblasts have an average CTSG F_ST_ of 0.185, which exceeds 80.5% of all random genes’ F_ST_. Hematopoietic cells, APCs, red blood cells, and natural killer cells were not significantly different from the 1000 random genes’ F_ST_. Other cell types had lower F_ST_ than the average for 1000 random genes, including eosinophils in the intestine and thrombocytes in the head kidney, suggesting that the CTSGs for these cells experience stabilizing selection that slows evolution. Note that natural killer cells had the highest gene-level F_ST_ values, but as this cluster had few CTSGs (3 loci), we are not confident these represent exceptional selection at the cell level.

**Figure 3.**
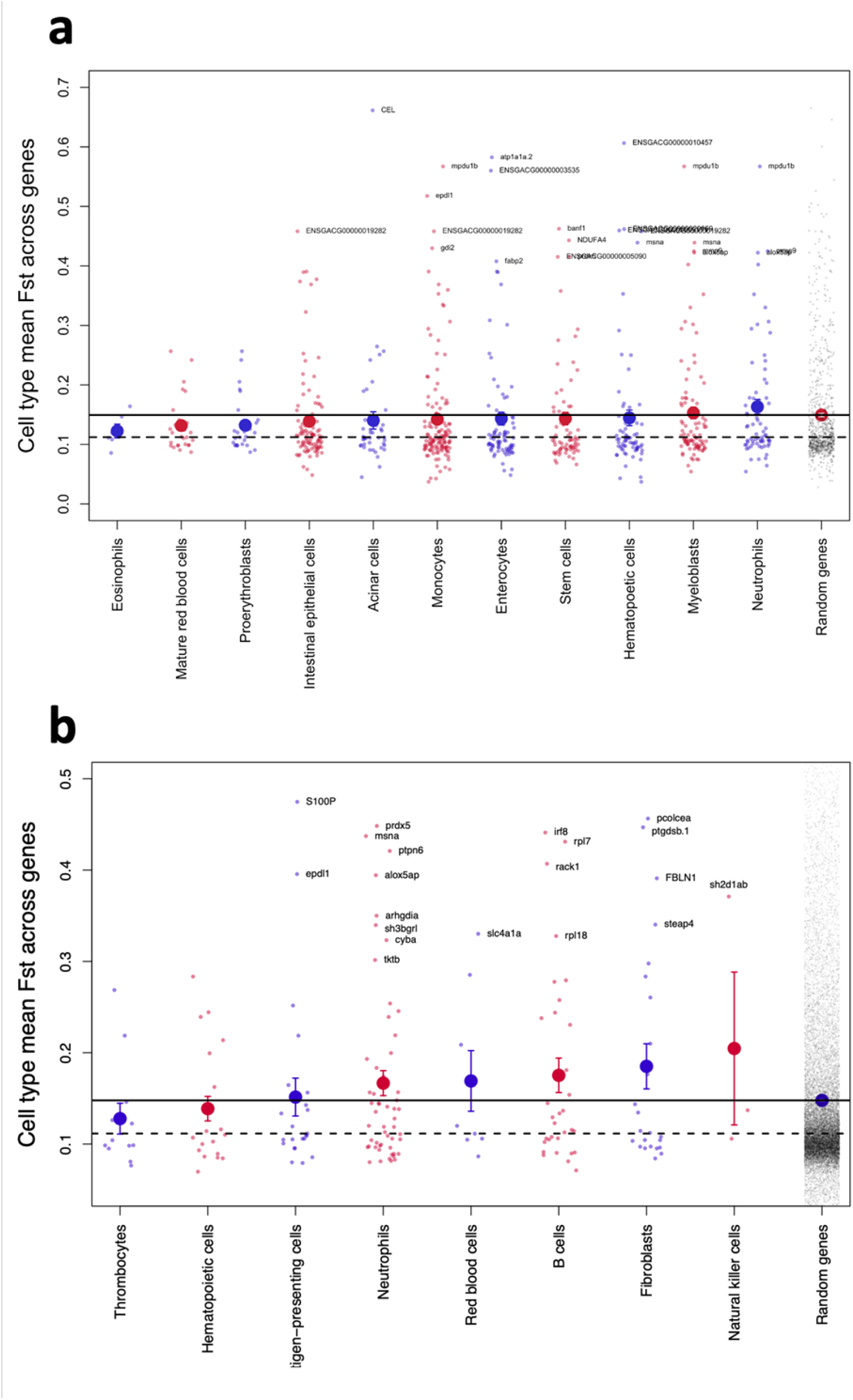
Evolutionary divergence between marine and freshwater stickleback (mean Fst) differs significantly between cell types as defined by **a)** intestinal single-cell RNAseq cell clusters, and **b)** head kidney scRNAseq cell clusters. Small data points are the cell-type-specific genes (CTSGs) that diagnose each cell type. Larger circles are averages across all CTSGs for a given cell type (with 1 standard error confidence intervals). Cell types are alternately colored. At the far right of each plot are divergence measures for 1000 coding genes sampled randomly from the genome (tiny black points) with the mean (solid dot and solid horizontal reference line) and median (horizontal reference line) for comparison. For each panel the top CTSGs (highest mean Fst) are labeled with gene names. Supplementary Figure 4 provides the equivalent figure with error bars for each gene.

Some noteworthy CTSGs exhibited particularly strong selection, even within cell types that otherwise were unremarkable on average. The strongest selection was observed for *cel* (a highly polymorphic gene in humans) within the acinar cell cluster (Figure 3a)^9^. This gene is involved in producing the enzyme carboxyl ester lipase, which is responsible for hydrolyzing dietary fat, esters, and some vitamins, and may be important in adaptation to changing nutrition during the transition from marine to freshwater habitat, as has been observed for the fatty acid desaturase gene *fads2*^10,11^.Other genes with strong selection in the intestine include ENSGACG00000010457 (currently undescribed) and *atp1a1a*.*2* (ATPase Na+/K+ transporting subunit alpha 1a, tandem duplicate 2). In the head kidney, we observed strong selection on *prdx5* in neutrophils, *pcolcea* in fibroblasts, and *s100p* in antigen-presenting cells (Figure 3b). S100P is present in mammals and teleosts, but is only partially represented in teleosts and was apparently lost in some teleost sublinages^12^. This gene is differentially expressed in stickleback in response to parasite infection or antigen injection^13^.

Both empirical and theoretical evidence suggest that genetic interactions may influence the evolutionary rate of genes, e.g., highly connected genes in a co-expression network may be evolutionarily constrained due to their greater pleiotropy^14,15^. To evaluate whether among-gene variation in evolutionary rate is influenced by gene network connectivity, we mapped loci included in this study onto a co-expression network. Briefly, using scRNA transcriptomic data from both anterior intestines and posterior intestines we reconstructed a coexpression network and calculated degree, eigenvector, Katz, betweenness and closeness centrality scores for each gene in the network. Using likelihood ratio comparisons of nested models with a fixed effect of immune-gene-status versus just an intercept we find significant variation in gene network centrality exists between immune- and non-immune-associated genes (LLR = 3.898, df = 1, P = 0.048). There is a significant positive correlation between closeness centrality and gene F_ST_ (Fig. 4a). Additionally, we observed that gene network centralities varied among scRNA-seq-defined cell clusters (Fig. 4b). However, the observed difference between immune- and non-immune-associated gene network centralities does not significantly explain the observed differences in F_ST._ (LLR = 0.410, df = 1, P = 0.522). Relaxing assumptions of linearity, we constructed a series of machine learning models–which allow for complex, non-linear relationships–to predict F_ST_ from gene network centrality measures. Based on cross-validation prediction performance, the machine learning models show at best a weak association between centrality and F_ST_ (Fig. 4c). Taken together, our analyses show that–despite correlations existing between gene network features and F_ST_–the network features alone are inadequate to explain observed variation in evolutionary rate.

**Figure 4:**
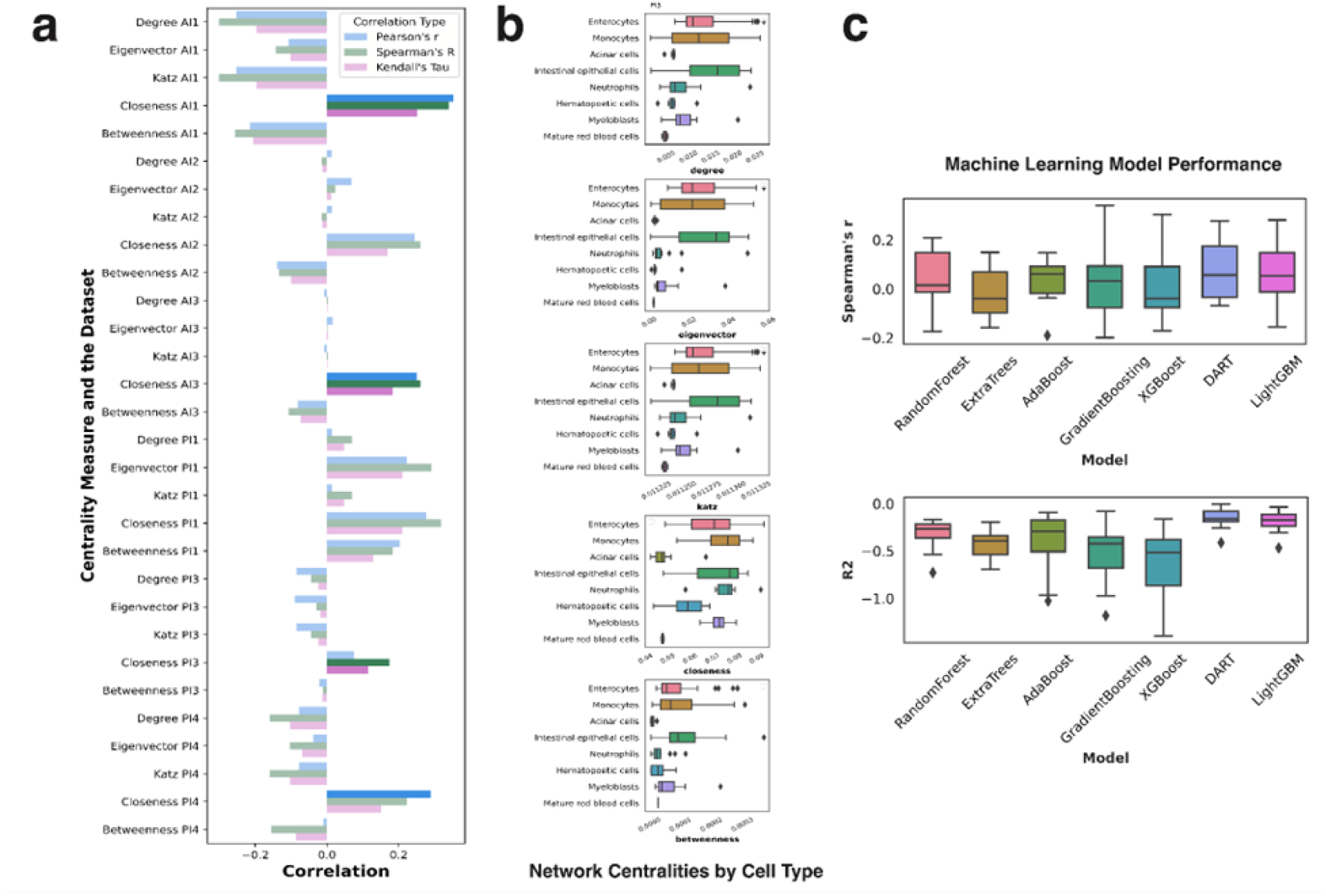
**a.** Correlations between individual network features and gene evolutionary rates. Significant correlations with p-values smaller than 0.05 were highlighted. **b.** Gene network centrality variations across cell types. **c.** Model performance summary by R2 score and Spearman’s rank correlation coefficient between the predicted gene F_ST_ and the true F_ST_ values, obtained through 5-fold cross-validation.

Biologists often expect that rates of molecular evolution on short time scales can be extrapolated to longer time scales. However, these time scales can be decoupled. For instance, strong frequency-dependent selection could lead to rapid fluctuations in allele frequencies over a few generations (and population divergence) that may not produce net genetic change over many generations. Therefore, in addition to our short-term measure of selection (F_ST_), we calculated metrics of selection over longer time scales (phylogenetic branch lengths and dN/dS). We first filtered the *G. aculeatus* genome annotation to remove all but the longest isoform for each gene using AGAT v1.0^16^. The remaining coding sequences were used to identify one-to-one orthologs (using Orthofinder v2.5.4) in the three other Gasterostiformes with whole genome species: *G. wheatlandi, Apeltes quadracus*, and *Aulorhynchus flavidus*^17^. Each gene alignment was trimmed using MACSE v2.07 and TrimAl v1.4.1 to remove non-homologous regions and regions with little data across the four species^18,19^. Trimmed gene alignments were then used to calculate dN/dS and tree length (all sites) using the CODEML program in PAML v4.9^20^. We then tested whether dN/dS and tree length for CTSGs differed between cell types, within each of four tissues (intestine, head kidney, gill, and liver).

The metric dN/dS measures the ratio of non-synonymous substitutions that might be visible to selection, to synonymous substitutions that are largely expected to be neutral. Stabilizing selection tends to lead to a low dN/dS when new mutations that change amino acid sequences tend to be harmful. In general, dN/dS for CTSGs are less than 0.5, implying overall that selective constraints tend to remove amino-acid-changing mutations (Figure 5). However, this ratio varied significantly among cell types (Fig. S1; liver F_10,188_ =2.69, P = 0.0042; intestine F_13,94_ = 5.15, P = 0.0024; head kidney F_5,177_ = 3.027, P = 0.0120; gills F_4,57_ = 1.854, P = 0.1320).

Within the liver, stem cells and eosinophils had the lowest dN/dS, whereas enterocytes, acinar cells, and erythroblasts had the highest dN/dS. In head kidneys, hematopoietic stem cells and natural killer cells had lower dN/dS than platelets or neutrophils. In the liver, hepatocytes had higher dN/dS than neutrophils, which were notably low. This raises an interesting observation that the CTSG for neutrophils in the liver are different from the CTSG for neutrophils in the head kidney, and the latter have higher dN/dS. Although there was no significant variation in dN/dS among cell types in gill samples, the trend is for natural killer cells and neutrophils to have lower dN/dS than B cells, epithelial cells, or erythrocytes. Also notably, dN/dS tended in general to be higher for CTSGs in intestinal cells (of any type) relative to CTSGs in the head kidney cells (Fig. S1a vs S1b).

We obtain somewhat similar results when we focus on phylogenetic tree length per gene as a metric of genic substitution rates. Genes under stronger positive selection tend to accumulate more substitutions overall resulting in greater overall branch lengths, whereas selective constraints can compress branch lengths. Cell types differed in tree length for cells in the liver (F_3,94_ =5.55, P = 0.0015), and head kidney (F_5,177_ =2.314, P = 0.0457), but not gill or intestine (both P>0.1), as illustrated in Supplementary Figure S4. In the head kidney, neutrophils and platelets again had higher evolutionary rates than hematopoietic stem cells. In the liver, hepatocytes had higher rates of substitutions than neutrophils (which were again lower than for neutrophils CTSGs in the head kidney).

Tree length and dN/dS were positively correlated (r = 0.489, t = 7.68, df = 188, P < 0.0001). However, across cell types, on average these macroevolutionary metrics were not correlated with microevolutionary rates (among-population F_ST_ versus dN/dS r = 0.317 P = 0.315; F_ST_ versus tree length r= -0.398 P = 0.200). This decoupling of microevolutionary rates (as measured by F_ST_) and macroevolutionary rates can have several causes. The most intriguing possibility is that frequency-dependent selection imposed by host-parasite coevolution can lead to rapid evolutionary cycling of alleles (contributing to rapid among-population divergence on short time scales), but long-term stasis as polymorphism is retained.

Rates of evolution (allele frequency change, substitution rates) are known to vary among genes within a genome. Some loci are subject to strong constraints that eliminate new mutations, whereas other loci experience persistent positive selection favoring the replacement of ancestral alleles by new mutations. This variation in evolutionary rate can be attributed to various genetic forces (varying mutation rates or recombination rates) and differences in the strength of selection. Immune genes in particular have been observed to evolve quickly, ostensibly because the antagonistic ‘arms race’ between hosts and pathogens or parasites leads to a constantly changing selection pressure. Analyses of these differences in rates of genic evolution are usually applied to genes individually, ignoring the joint action of genes. Or, analyses use coarsely defined Gene Ontology (GO) categories that are usually not validated for the species being studied (nor for any close relatives). A useful alternative that we explore here is to leverage data-driven functional roles, obtained from the same study organism using single-cell RNAseq data. The merger of scRNAseq data with population genomics allows gene function assignments (by cell cluster) within the study organism. We use this approach to show that the rate of genic evolution varies between scRNAseq-defined cell clusters, for intestinal cells, head kidney cells, liver, and gills. In these tissues, the faster evolving cell clusters are consistently central players in the adaptive and innate immune systems. We are thus able to identify cell types whose function is likely to evolve especially quickly, and conserved cell types. Within a given cell type we can also identify the cell-type-specific genes that are evolving especially quickly (e.g., *polcea* in fibroblasts). This merger of scRNAseq and population genomic approaches thus provides a distinctive window into functional interpretation of evolutionary genomic data, and lends further support to the insight that immune function is an especially labile trait over microevolutionary time.

## Supporting information

Captions for Supplemental Figures and Tables

## Data and code repositories

Poolseq population genomic data are archived on the SRA (accession numbers listed in Weber et al 2022). Single cell RNAseq data are currently being archived. R code to generate statistical results and graphics, and associated intermediate data files (F_ST_, dN/dS) with metadata are currently being archived with FigShare.

## Acknowledgements

We gratefully acknowledge the contribution of JAX-UConn Single Cell Genomics Center, comprising the Single Cell Biology service, Genome Technologies and Cyberinfrastructure high performance computing resources at The Jackson Laboratory, for expert assistance with the work described herein. These shared services are supported in part by the JAX Cancer Center (P30 CA034196). We specifically acknowledge William Flynn and Martine Seignon for bioinformatic assistance. This work was supported by grants from the NIH (NIAID 1R01AI123659-01A1) and the Gordon and Betty Moore Foundation (GBMF9323 to DIB and KMM)

## Author contributions

The study was conceived and designed by DIB and MLR. Single cell sequencing and analyses performed by MLR, LEF, and MS. Population genomic data collection and analyses were conducted by AR and SS. Macroevolutionary rate analyses by DJ. Network analyses by WH and SVS. Statistical analyses and graphics by DIB and MLR. Funding acquisition DIB, KMM, NCS, and RC. Initial draft by MLR, DIB, WH, DJ, and MS with comments and feedback from SS, LEF, SVS, AR, KMM, NCS, and RC.

## Methods

### Single cell RNA sequencing

Single cell libraries were generated from the intestines of three individual laboratory-reared threespine stickleback. The parents of these laboratory-reared fish were from Sayward Estuary in British Columbia, Canada. For head kidneys, single cell libraries were generated from nine total laboratory-reared threespine stickleback. Three of these fish had parents from Sayward Estuary (as described for the intestines), three had parents from Roberts Lake and three had parents from Gosling Lake (all in British Columbia, Canada). To obtain fertilized eggs, gravid female fish from each source (Sayward Estuary, Roberts Lake, and Gosling Lake) were stripped of eggs and these eggs were mixed with sperm from macerated testes, taken from male stickleback in the same location as the female (Sayward Estuary, Roberts Lake, or Gosling Lake). All fish were collected in accordance with the Ministry of Forests, Lands, and Natural Resource Operations of British Columbia (Scientific Fish Collection permits NA12-77018 and NA12-84188). Fertilized eggs were transported to the University of Texas at Austin where fish were raised for two years before being transferred to the University of Connecticut.

To generate cell suspensions, stickleback were euthanized with MS-222 (IACUC: A21-025) and intestines or head kidneys were removed. Upon removal, intestines were initially cut into two pieces (each representing half of the total length of the intestine; these pieces were marked as the anterior and posterior intestine). The head kidneys were left intact. Each piece of the intestine or whole head kidney was placed into a sterile 24 well plate on ice with 2 mL on R-90 media (90% (v/v) RPMI 1640 (+L-glutamine, -Phenol red; Gibco) + 10% (v/v) distilled water) in each well. From there, each piece of the intestine was cut longitudinally and then sliced into small fragments; head kidneys, gills, and livers remained intact. The intestinal pieces and whole head kidneys, livers, and gills were dissociated physically by using a sterile pipette tip. Liquid from each well of the plate was then individually passed through a 40um nylon screen (BD Falcon) and 2 mL of chilled R-90 was added to the resulting cell suspension. The suspension was spun for 10 min at 4°C and 550 *g*. Supernatant was discarded from each tube and cells were resuspended in 2 mL of chilled R-90. Once again, the suspension was spun for 10 min at 4°C and 550 *g*. Supernatant was discarded from each tube, and cells were resuspended in 1 mL of chilled R-90. Cells were then transferred on ice to The Jackson Laboratory for Genomic Medicine in Farmington, CT for sequencing.

### Single Cell Library Preparation and Sequencing

Once cells were transferred to The Jackson Laboratory for Genomic Medicine (within 1 hour of resuspension), cells were washed and suspended in PBS containing 0.04% BSA and immediately processed as follows. Cell viability was assessed on a Countess II automated cell counter (ThermoFisher), and an estimated 12,000 cells were loaded onto one lane of a 10x Genomics Chromium Controller. Single cell capture, barcoding, and single-indexed library preparation were performed using the 10x Genomics 3’ Gene Expression platform, v3.1 chemistry, and according to the manufacturer’s protocol (#CG000204)^21^. cDNA and libraries were checked for quality on Agilent 4200 Tapestation, quantified by KAPA qPCR, and sequenced on an Illumina NovaSeq 6000 targeting 6,000 barcoded cells with an average sequencing depth of 50,000 read pairs per cell (approximately 13% of an S4 flow cell).

*scRNA-seq Data Processing and Analysis*. Illumina base call files for all libraries were converted to FASTQs using bcl2fastq v2.20.0.422 (Illumina) and FASTQ files were aligned to reference genome constructed from the 2020v5 *G. aculeatus* assembly and annotation files available at https://stickleback.genetics.uga.edu/^22^. Briefly, annotations from Ensembl (release 95) were combined with repeat, Y chromosome, and revised annotations from Nath et al. using AGAT (0.4.0) and a STAR-compatible reference genome was generated Cell Ranger (version 6.0.0, 10x Genomics) using these annotations and the v5 assembly from Nath et al^23^. The Cell Ranger count pipeline was used to construct cell-by-gene counts matrix for each library, subsequently analyzed using Scanpy 1.3.7 and the Loupe Cell Browser (10x Genomics)^24^.

Each counts matrix was individually subjected to quality control filtering, such that cells with fewer than 400 genes, more than 20% mtRNA content, and more than 500 hemoglobin transcripts were discarded from downstream analysis. Additionally, cells with more than 12,500, 15,000, and 25,000 UMI counts were discarded from libraries DB21009, DB2012, and DB21007, 008, 010, 011, respectively. Filtered counts matrices were concatenated, normalized by per-cell library size, and log transformed. The expression profiles of each cell at the 3,500 most highly variable genes (as measured by dispersion^21,25^) were used for principal component (PC) analysis and subsequently batch corrected using Harmony^26^. The batch corrected PCs were utilized for neighborhood graph generation (using 40 nearest-neighbors) and dimensionality reduction with UMAP^27^. Clustering was performed on this neighborhood graph using the Leiden community detection algorithm^28^. Subclustering was performed on a per-cluster ad hoc basis to separate visually distinct subpopulations of cells. This UMAP embedding and clustering metadata were then imported into the Loupe Cell Browser (using Cell Ranger aggr, version 6.0.0) for interactive analysis.

### Pool-seq

We used unbaited minnow traps to collect threespine stickleback from 15 lakes and one anadramous marine population (Sayward Estuary) in British Columbia (Table 1). Collections from these lakes were approved by University of Texas IACUC (07-032201) and a Scientific Fish Collection Permit from the Ministry of the Environment of British Columbia (NA07-32612). In addition we trapped stickleback from eight lakes in Alaska (IACUC protocol UConn A19-015, Alaska DFG collecting permit SF2019-085). We obtained fin clips from 100 fish per population, and extracted DNA using magnetic beads modifying the DNA extraction protocol by Bio-On-Magnetic-Beads^29^. Modifications included different lysis buffer (10 mM Tris pH 8.3, 50 mM KCl, 1.5 mM MgCl2, 0.3% Tween-20, and 0.3% NP-40). We added 255 μl of the lysis buffer and 20 μl of Proteinase K per sample, left overnight at 56 C. We also shaked the plate for 6 min with the isopropanol, added 120 ul of beads, and added 300 ul of isopropanol for the first wash. We did three 80% ethanol washes, all as indicated in the protocol. For the final elution we shake the plate for 7 min. We used modified Serapure beads for the extractions. We quantified DNA extract concentrations using picogreen on a Tecan-Magellan plate flourometer, and used these concentrations to pool equimolar amounts of DNA from each of 100 fish per population to generate one DNA pool per population. These pools were subjected to whole genome sequencing at a target depth of 200x (e.g., 1x per allelic copy), using a Illumina NovaSeq 6000 at the Center for Genome Innovation in the University of Connecticut. In addition to our own sample sequences we obtained poolSeq data from an Alaskan marine population (Rabbit Slough) and three additional Alaskan lake populations recently founded by marine fish^30^.

The 28 populations included 24 newly sequenced populations and 4 populations with sequence downloaded from NCBI SRA database (PRJNA671824)^31^. The raw sequencing reads for the 24 newly sequenced population are deposited in NCBI SRA Bioproject PRJNA850376. The number of individuals included in each population pool is in Supplementary Table 4. The populations were sequenced to an average coverage of 100-150, which is sufficient to allow an accurate estimation of allele frequencies. We used the pp_align program in the PoolParty pipeline to process the raw reads and to call SNP variants^32^. The parameter settings we used in pp_align was BQUAL=20, MAPQ=10, SNPQ=20, MINLENGTH=25, INWIN=2, MAF=0.00005, MINDP=10. Then, we used the R package poolfstat to calculate the pairwise Fst values for each SNP, using the following settings: min.rc=4, min.cov.per.pool = 10, max.cov.per.pool = 2000, nlines.per.readblock = 1e+06,noindel=T^33^. We used the R package matrixStats to calculate the mean and standard deviation of the Fst values across all pairwise comparisons of freshwater populations, freshwater populations within each region, anadromous Rabbit Slough and Alaska freshwater populations, anadromous Sayward and British Columbia freshwater populations, and anadromous Rabbit Slough and recently freshwater Alaskan populations Scout, Loberg, and Cheney^34^. We then used the R package dplyr to filter the SNPs and associated Fst data by the base pair locations of genes of interest identified through scRNA, with 5k base pairs added on either side of the location^35^. 1000 random genes were sampled from all stickleback genes using the R function sample, and dplyr was used to filter the SNPs and associated Fst data by the base pair locations of the random genes with 5k base pairs added on either side.

### Network Analysis

There were six sets of intestine scRNA-seq data used for generating single-cell gene co-expression networks. These datasets consist of three anterior intestine scRNA-seq datasets (AI1, AI2, AI3) and three posterior intestine scRNA-seq datasets (PI1, PI3, PI4). For each scRNA-seq dataset, we generated a gene co-expression network by calculating pairwise Spearman’s rank correlation coefficient among the gene pairs. Fully-connected weighted co-expression networks were used for network feature extraction. Cells and genes with close-to-zero variation were filtered out. Specifically, for AI1, AI2, AI3, PI1, PI3, and PI4, 20%, 10%, 0%, 10%, 10%, and 10% of the cells, respectively, and 36%, 36%, 46%, 36%, 47%, and 35% of the genes with the least variance after scaling were removed. Nodes in these networks represent genes and the edge weight between any pair of two genes represents the similarity of their expression profile across samples in the intestine single-cell dataset. For each gene, we considered degree centrality, eigenvector centrality, Katz centrality, and closeness centrality. A range of regression models were evaluated for their performance in predicting gene F_ST_ when the extracted network features were used as predictors.

## Notes

### Competing Interest Statement

The authors have declared no competing interest.

