## Supplementary material for "Rates of evolution differ between cell types identified by single-cell RNAseq in threespine stickleback": Captions for Supplemental Figures and Tables

**Figure S1.** The cell-type specific genes of various cell populations differ in the strength of purifying selection over macroevolutionary time. We measure selection by calculating dN/dS ratios between stickleback species. dN/dS is the ratio of non-synonymous to synonymous substitutions in coding sequences, and is smaller for genes subject to stronger purifying selection that slows evolution. We present dN/dS for cell-type specific genes, for each cell type within each of four tissues. With the exception of gills, all tissue types exhibit statistically significant differences in dN/dS between cell types.

**Figure S2.** The frequency distribution of marine versus freshwater Fst values for SNPs in **A)** 1000 randomly selected genes (black), and in **B)** the cell-type specific genes (purple). These distributions have equivalent means (Fst = 0.1463 and 0.1468 respectively, $t = -0.839$, $P = 0.4013$, df = 130,276), but slightly different distributions (KS-test $D = 0.0085$, $P = 0.0005$) due to greater variance in the CTSG SNPs (0.0228 vs 0.0215, F-test $P < 0.001$).

**Figure S3.** Variation in allele frequency divergence (Fst) among cell-type specific genes. Each cell-type specific gene is represented along the x axis. Each SNP within each gene (including 5kb of flanking sequence) is represented by a point, which is the mean Fst between marine and freshwater populations for that given SNP. Averaging across SNPs within a gene yields the mean Fst for an entire gene (blue points). Genes are ranked by mean Fst, so genes on the right exhibit the strongest signal of divergent selection.

**Figure S4.** As in Figure 3, but replotted with genic tree length rather than dN/dS.

**Table S1.** Identified clusters from the intestine with all locally distinguishing significant genes.

**Table S2.** Identified clusters from the liver with all locally distinguishing significant genes.

**Table S3.** Identified clusters from the gills with all locally distinguishing significant genes.
